# Audiovisual gamma stimulation enhances hippocampal neurogenesis and neural circuit plasticity in aging mice

**DOI:** 10.1101/2025.01.13.632794

**Authors:** Mariela F. Trinchero, Magalí Herrero, Matías Mugnaini, Andrea Aguilar-Arredondo, Sabrina Benas, Ignacio G. Satorre, Emilio Kropff, Alejandro F. Schinder

**Author notes:** These authors contributed equally to this work (Mariela F. Trinchero, Magalí Herrero).

## Abstract

Gamma oscillations are disrupted in various neurological disorders, including Alzheimer’s disease (AD). In AD mouse models, non-invasive audiovisual stimulation (AuViS) at 40 Hz enhances gamma oscillations, clears amyloid-beta, and improves cognition. We investigated mechanisms of circuit remodeling underlying these restorative effects by leveraging the sensitivity of hippocampal neurogenesis to activity in middle-aged wild-type mice. AuViS increased progenitor cell proliferation, neuronal differentiation and morphological maturation of newborn granule cells, promoting their synaptic integration. While visual or auditory stimuli alone induced dendritic growth, axonal changes required combined audiovisual stimulation. The actions of AuViS involved neurotrophin pathways, as shown by the lack of effect upon TrkB signaling blockade. These results reveal widespread plasticity mechanisms triggered by AuViS, a therapeutic approach currently proposed for treating neurological disorders in humans.

## Introduction

Understanding the fundamental mechanisms of brain senescence is a critical problem in neuroscience since aging represents the primary risk factor for degeneration of the nervous system. The cognitive decline associated with physiological aging and neurodegeneration often results from impaired synaptic transmission and plasticity, which correlates with disruptions in gamma brain oscillations (*1*). Recent work using optogenetic activation of parvalbumin-expressing GABAergic interneurons has shown that stimuli that enhance the gamma power at 40 Hz can improve inflammatory responses and reduce the levels of amyloid beta (Aβ) peptides in mouse models of Alzheimer’s disease (AD) (*2*). Remarkably, non-invasive 40 Hz audiovisual stimulation (AuViS) can also enhance gamma oscillations in several brain regions and exert neuroprotective effects, including prevention of memory impairment and reduction in the levels of Aβ peptides (*2, 3*). AuViS has also been beneficial in models of stroke, enhancing long-term potentiation and upregulating genes related to plasticity, while reducing cell death (*4*). Known consequences of 40 Hz AuViS include changes in transcriptomic profiles, morphological modifications in microglia, increased Aβ engulfment, and improved glymphatic clearance, suggesting anti-inflammatory mechanisms (*5*). In addition to its anti-inflammatory effects, gamma stimulation by AuViS may trigger unique mechanisms of circuit remodeling that enhance cognitive function, as suggested by the improved memory performance observed in animal models of neurodegenerative disorders. The safety, affordability, and simplicity of 40 Hz AuViS have prompted clinical trials investigating its efficacy across various pathologies, which underscores the need for a thorough understanding of how such stimulation affects neuroplasticity (*5*).

One of the most striking forms of structural plasticity is neurogenesis in the dentate gyrus of the adult hippocampus. This process involves the proliferation and differentiation of neural progenitor cells and the subsequent integration of adult-born granule cells (aGCs) into the existing circuits (*6–8*). Aging reduces the neurogenic capacity of neural stem cells, delays the differentiation and integration of aGCs, and impairs their survival (*9, 10*). However, exposing aging mice to exercise and environmental enrichment (EE), which increase neuronal activity in the dentate gyrus, significantly enhance aGC differentiation and integration into the local circuit (*11–14*). We have hypothesized that neurogenesis in the aging brain could serve as a reliable probe for investigating additional mechanisms underlying the restorative effects of AuViS. We found that AuViS enhances activity in the dentate gyrus and increases proliferation and neuronal differentiation of progenitor cells. AuViS at 40 Hz induced the growth of aGCs as well as their pre- and postsynaptic integration. However, visual or auditory stimuli alone promoted dendritic growth but failed to induce presynaptic changes. The effects of AuViS engaged neurotrophin signaling, a well-known activity-dependent mechanism for circuit remodeling. These results highlight the potential of multisensory gamma stimulation as a putative therapeutic approach for neurodegenerative diseases and age-related cognitive decline.

## Results

### AuViS activates the aging dentate gyrus and triggers the maturation of newborn aGCs

We sought to leverage the sensitivity of neurogenesis in middle-aged mice to interrogate the effects of AuViS on circuit plasticity. To investigate the extent to which 40 Hz non-invasive sensory stimulation affects the activity of the dentate gyrus, we assessed the expression of the immediate early gene Arc in response to AuViS in 8-month-old (8M) mice (**Fig. 1A**). Animals were exposed to 40 Hz AuViS (**Fig. S1**) or lights off (control) for 2 h, and Arc expression in the granule cell layer was assessed 90 min later. Compared to control conditions, AuViS induced a 2-fold increase in the proportion of Arc/NeuN^+^ cells in the subgranular zone, but not in other areas of the granule cell layer (**Fig. 1B,C**). This result suggests that AuViS preferentially recruits neurons within the subgranular zone, the area where neural progenitor cells and immature aGCs are located. We therefore tested the effects of AuViS on developing aGCs from 8M mice, labeled by retroviral expression of GFP. Animals were exposed to AuViS (40 Hz, 4h/day) after retroviral delivery, and GFP^+^-aGCs were analyzed 17 days post infection (dpi) by confocal imaging to assess the morphological architecture of their dendritic tree (**Fig. 1D**). Newborn aGCs from mice exposed to AuViS displayed a 2-fold increase in their dendritic length and branching points, indicating a more advanced state of maturation compared to the control group (**Fig. 1E,F**). GCs form glutamatergic connections with CA3 pyramidal cells via mossy fiber boutons (MFBs) and recruit feedforward GABAergic inhibition through filopodial extensions (*15–19*) (**Fig. 1G**). AuViS elicited a substantial increase of both MFB area and filopodial number, indicating an enhanced recruitment of both output target cells (**Fig. 1H,I**). In order to determine if the observed effects were produced by AuViS at 40 Hz or solely due to stimulation by light and sound, mice were exposed to audiovisual flicker maintaining the 50% working cycle but using random frequency. Random stimulation failed to induce structural modifications at the pre- or postsynaptic levels (**Fig. 1J-L**). Overall, we found that 40 Hz stimulation promotes dendritic growth, suggesting faster innervation by afferent axons from the entorhinal cortex, along with a more mature morphological development of synapses with pyramidal cells and GABAergic interneurons in CA3.

**Fig. 1.**
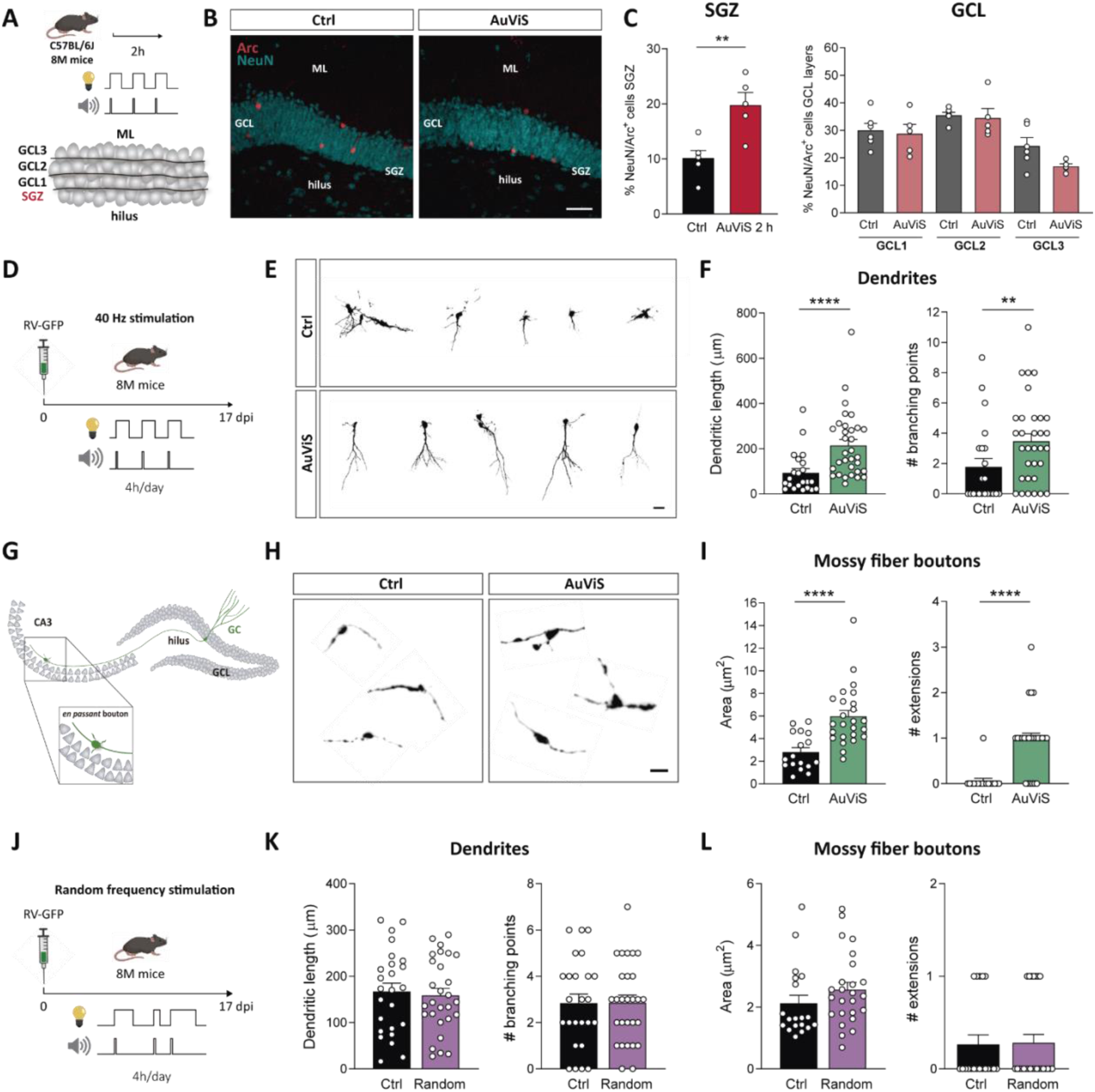
40 Hz AuViS promotes structural remodeling of new aGCs in middle-aged mice. (**A-C**) Acute effects of AuViS on the activation of the granule cell layer (GCL). (**A**) Top: Experimental design. 8M mice were exposed to 40 Hz AuViS for 2 h. Bottom: Schematic diagram depicting four subdivisions of the GCL. (**B**) Representative images of Arc expression (red) in GCL labeled with NeuN (blue). ML: molecular layer, SGZ: subgranular zone. Scale bar, 50 μm. (**C**) Quantification of Arc expression in the SGZ and GCL. (**) denotes *p*<0.01 after Mann–Whitney test. Sample size (slices/mice): 6/4 (Ctrl) and 5/4 (AuViS). (**D–I**) Effects of chronic 40 Hz AuViS on aGC development. (**D**) RV-GFP injection was followed by 17 days of 40 Hz AuViS. (**E**) Representative confocal images of 17-dpi GFP-aGCs from control (upper panel) and stimulated (lower panel) mice. Individual neurons have been cropped from original images. Scale bar, 20 μm. (**F**) Dendritic measurements of 17-dpi GFP-aGCs. (**) and (****) denote *p*<0.01 and *p*<0.0001 after Mann– Whitney test. Sample sizes (neurons/mice): 22/3 (Ctrl) and 32/4 (AuViS). (**G**) Scheme depicting the CA3 location of MFBs under study. (**H**) Representative confocal images of MFBs from 17 dpi aGCs for both groups. Scale bar, 5 μm. (**I**) Quantitative analysis of MFB morphology. (****) denotes *p*<0.0001 after Mann–Whitney test. Sample sizes (neurons/mice): 17/3 (Ctrl) and 25/4 (AuViS). (**J-L**) Effects of random AuViS on aGC development. (**J**) RV-GFP injection was followed by 17 days of exposure to random AuViS. (**K**) Dendritic measurements of 17-dpi GFP-aGCs. Sample sizes (neurons/mice): 25/4 (Ctrl) and 28/4 (Random). (**L**) MFB measurements. Sample sizes (neurons/mice): 19/4 (Ctrl) and 25/4 (Random). Bars denote mean ± SEM.

Given the effects observed on aGCs with combined auditory and visual stimulation, we sought to further investigate the individual contributions of each sensory modality. First, we examined the impact of visual flicker on the modulation of extracellular local field potentials (LFPs) recorded through chronic tetrode implants in freely moving mice (**Fig. 2A**). All implanted animals exhibited at least one tetrode located within or in close proximity to the granule cell layer, where the LFP displayed modulation at 40 Hz (**Fig. 2B-D and S2**). Forty Hz visual flickering acutely activates local networks of the dentate gyrus enhancing the amplitude of the 40 Hz component of gamma oscillations. Next, we examined the dendritic and axonal projections of 17 dpi GFP^+^ aGCs born in 8M mice exposed to 4h/day of visual stimulation at 40 Hz (**Fig. 2E**). This analysis revealed a 2-fold increase in the dendritic length compared to the control group, similar to the effects exerted by AuViS (**Fig. 2F,H**). However, we found no changes in the area of MFBs or their filopodial extensions (**Fig. 2G,I**), suggesting that visual input selectively enhances dendritic development without affecting the structural properties of the presynaptic terminals in developing aGCs. The effect of visual flicker was remarkably solid, since reducing the stimulation window to 1 h per day still exerted considerable trophic effects on the overall dendritic architecture in 21-day-old aGCs (**Fig. S3**). In contrast, visual stimulation at 8 Hz did not elicit morphological changes, which points to the idea that the trophic effects described above are specific to frequencies within the gamma range (**Fig. S4**).

**Fig. 2.**
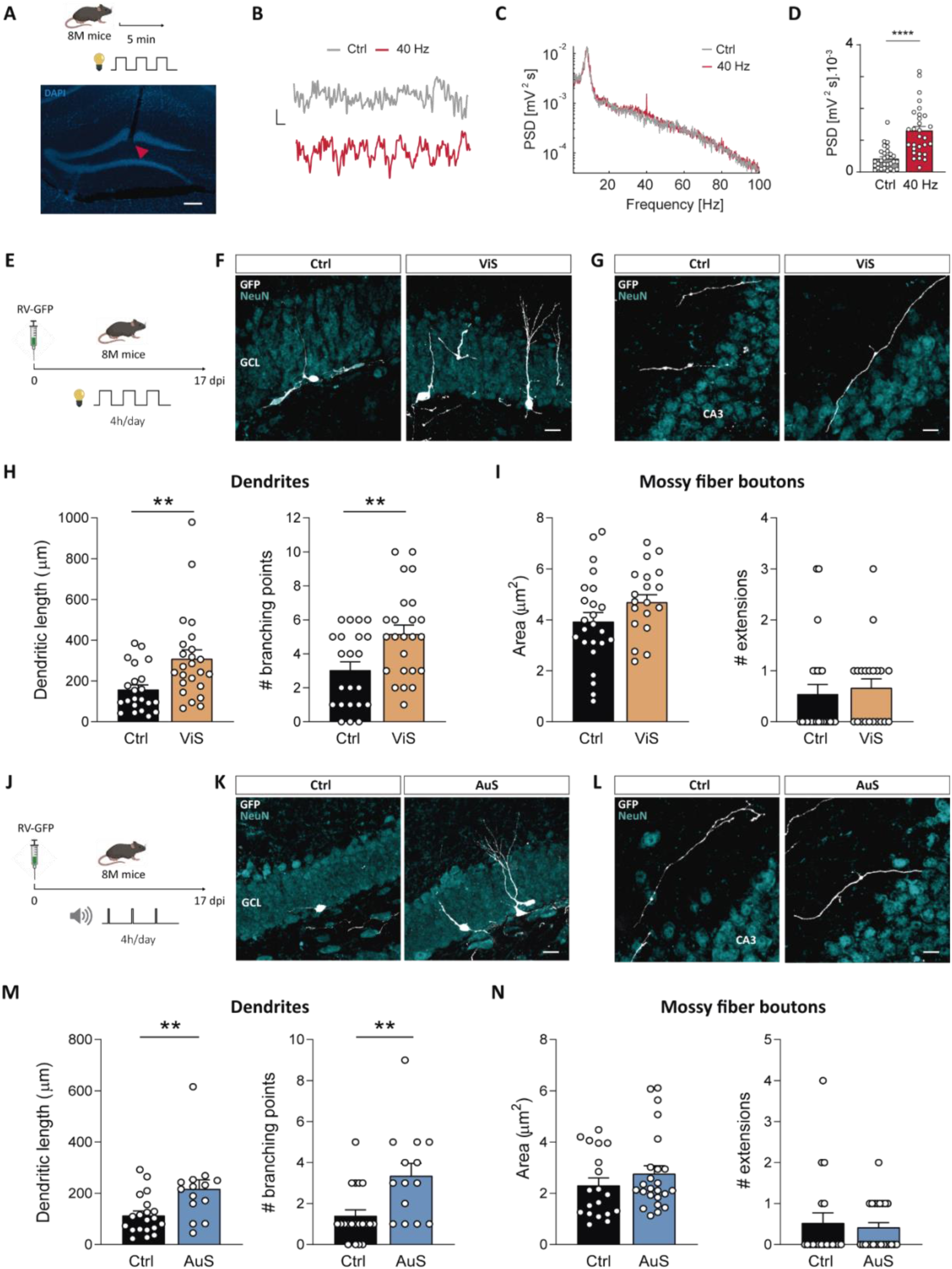
Dendritic remodeling by visual or auditory stimuli. (**A-D**) Acute effects of light flicker (ViS) on the activation of the dentate gyrus by in vivo recordings. (**A**) Top: Experimental design. Mice were exposed to 5 min of 40 Hz ViS or lights-off. Bottom: Histology showing the trace of a tetrode targeted to the granule cell layer (red arrowhead: tetrode tip). Scale bar, 50 μm. (**B**) Representative example trace of 1 s raw local field potential during flickering (40 Hz) or lights-off (Ctrl) conditions. Scale bars, 1 mV, 0.1 s. (**C**) Corresponding power spectral density for 5 min of flickering (40 Hz) or lights-off (Ctrl). (**D**) Distribution of the power spectral density at 40 Hz for consecutive non-overlapping 10 s windows of data. (****) denotes *p*<0.0001 after Mann– Whitney test. (**E-I**) Effects of 40 Hz light flicker. (**E**) 8M mice were injected with a RV-GFP in the dentate gyrus and exposed to 17 days of 40 Hz ViS 4h/day. Morphological parameters were analyzed in 17 dpi GFP-aGCs by confocal imaging. (**F**) Representative confocal images (GFP: white, NeuN: blue). Scale bar, 20 μm. (**G**) Representative images of MFBs in CA3. Scale bar, 20 μm. (**H**) Dendritic measurements. (**) denotes *p*<0.01 after Mann–Whitney test. Sample sizes (neurons/mice): 22/4 (Ctrl) and 24/4 (ViS). (**I**) MFB morphology. Sample sizes (neurons/mice): 24/4 (Ctrl) and 21/4 (ViS). (**J-N**) Effects of 40 Hz sound clicks (AuS) on 17-dpi GFP-aGCs. (**J**) RV-GFP injected mice were exposed AuS (4 h/day, 17 days). (**K**) Representative confocal images. Scale bar, 20 μm. (**L**) Representative MFBs. Scale bar, 20 μm. (**M**) Dendritic measurements. (**) denotes p<0.01 after Mann–Whitney test. Sample sizes (neurons/mice): 20/3 (Ctrl) and 14/3 (AuS). (**N**) MFB morphology. Sample sizes (neurons/mice): 19/3 (Ctrl) and 24/3 (AuS). Bars denote mean ± SEM.

To examine the specific contribution of auditory stimulation, mice were exposed to 10 kHz pulses delivered at 40 Hz 4h/day to analyze the morphology of GFP^+^ aGCs born in 8M animals at 17 dpi (**Fig. 2J**). Sound stimulation induced a significant increase in dendritic length and complexity with no presynaptic modulation, reflecting a trophic action that was similar to the one observed with visual stimulation alone (**Fig. 2K-N)**. Together, these results reveal that combined audiovisual stimulation exerts a synergistic effect on presynaptic terminals of aGCs, which is crucial for promoting their complete integration into the circuit, an outcome that cannot be attained with either stimulus alone.

### AuViS promotes synaptic integration of newborn aGCs

The structural plasticity described above suggests that AuViS induces overall maturation of aGCs at multiple levels, including traits required for neuronal function and synaptic integration. To test this possibility, electrophysiological recordings were carried out in 17-day-old aGCs using 8M *Ascl1^CreERT2^;CAG^floxStoptdTomato^*mice. Upon tamoxifen injection, the red fluorescent reporter tdTomato was expressed in neural stem cells and their progeny, which allowed labeling a substantial number of new aGCs to perform whole-cell recordings. Neurons from mice exposed to AuViS exhibited a decrease in their input resistance, compatible with a higher density of ion channels required for the membrane excitability to become mature. However, aGCs from stimulated and control groups failed to develop sufficient excitability to fire action potentials (**Fig. S5**). Thus, we examined the effects of AuViS on the functional properties of 24 dpi aGCs, as they are closer to the onset of synaptic integration (**Fig. 3A**). TdTomato^+^-aGCs from mice exposed to AuViS displayed a more mature (hyperpolarized) resting potential and higher membrane capacitance, which denotes a larger cell area. No differences were observed in the input resistance or threshold current for spiking (**Fig. 3B-D**). However, aGCs from control mice fired only a single action potential, whereas those from the AuViS group exhibited reliable spiking, indicative of a more advanced maturation state (**Fig. 3C-E**). Finally, spontaneous excitatory postsynaptic currents (sEPSCs) were measured to assess synaptic integration. While neurons from the AuViS group displayed a 2-fold increase in sEPSC frequency, the sEPSC amplitude remained unchanged, indicating that AuViS induced the formation of glutamatergic contacts without modifications in synaptic strength (**Fig. 3F,G**). These results demonstrate that sensory stimulation at 40 Hz not only boosts the maturation of new aGCs but also facilitates their functional integration, which suggests an active participation of the preexisting circuits.

**Fig. 3.**
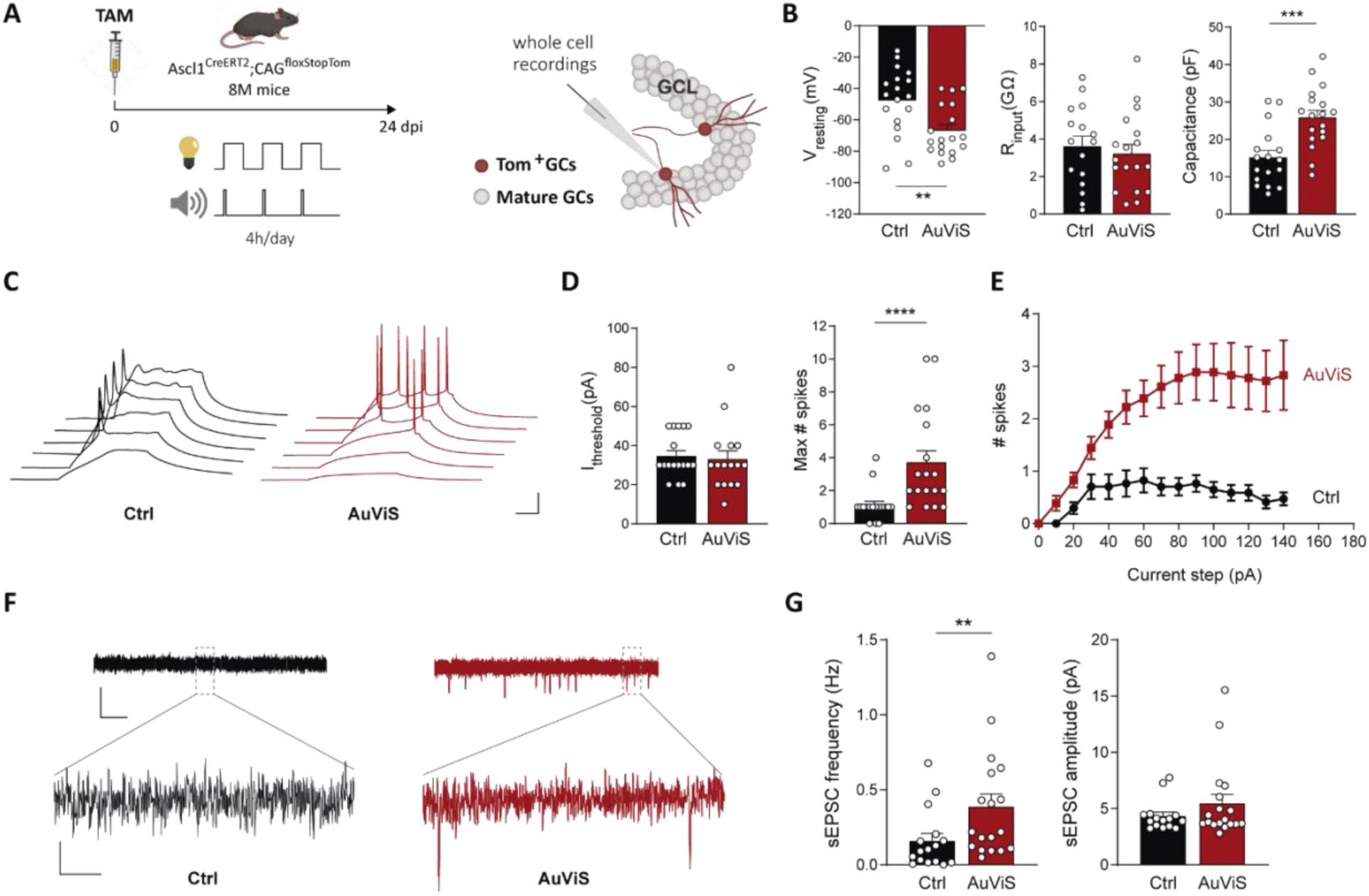
40 Hz AuViS triggers functional integration of aGCs. (**A**) Experimental design. Eight-month-old Ascl1^CreERT2^;CAG^floxStopTom^ mice received TAM to label aGCs. Acute slices were prepared 24 days after 40 Hz AuViS or control treatments to assess intrinsic properties and connectivity using whole-cell recordings in tdTomato^+^-aGCs. (**B**) Resting potential (left), input resistance (middle) and membrane capacitance (right). Sample sizes (neurons/mice): 17/3 (Ctrl) and 18/4 (AuViS). (**) and (***) denote *p*<0.01 and *p*<0.001 after Mann–Whitney test. (**C**) Representative voltage traces in response to depolarizing current steps (20–70 pA). Scales, 25 mV, 50 ms. (**D**) Threshold current for spiking (left), and maximum number of spikes elicited by step depolarizations (right). Sample sizes (neurons/mice): 17/3 (Ctrl) and 18/4 (AuViS). (****) denotes *p*<0.0001 after Mann–Whitney test. (**E**) Number of action potentials elicited by current injections (V_resting_ =-70 mV). Sample sizes (neurons/mice): 15/3 (Ctrl) and 15/4 (AuViS). (**F**) Representative voltage-clamp traces depicting sEPSCs (V_holding_ =-70 mV). Scale bars, 10 pA, 1 s (top); 1 pA, 25 ms (bottom). (**G**) Frequency and amplitude of sEPSCs measured during 120 s. (**) denotes *p*<0.01 after Mann–Whitney test. Sample sizes (neurons/mice): 16/3 (Ctrl) and 18/4 (AuViS). All data denote mean ± SEM.

### The actions of AuViS require neurotrophin signaling

Neurotrophins are critical for activity-dependent circuit plasticity from development to adulthood (Poo 2013 and Trinchero 2017). Given that 40 Hz stimulation enhances activity in the dentate gyrus, we hypothesized that AuViS might promote plasticity via TrkB signaling. Using a retrovirus to overexpress Lrig1, a negative regulator of TrkB neurotrophin receptors, we labeled aGCs from 8M mice exposed to AuViS and analyzed aGCs 17 days later (**Fig. 4A**) (*20–22*). In line with previous results, aGCs from mice exposed to AuViS displayed a 2-fold increase in dendritic length, compared to controls (**Fig. 4B,C**). Notably, oeLrig1 abolished AuViS-induced dendritic growth, revealing a critical role of neurotrophin signaling at the cell autonomous level.

**Fig. 4.**
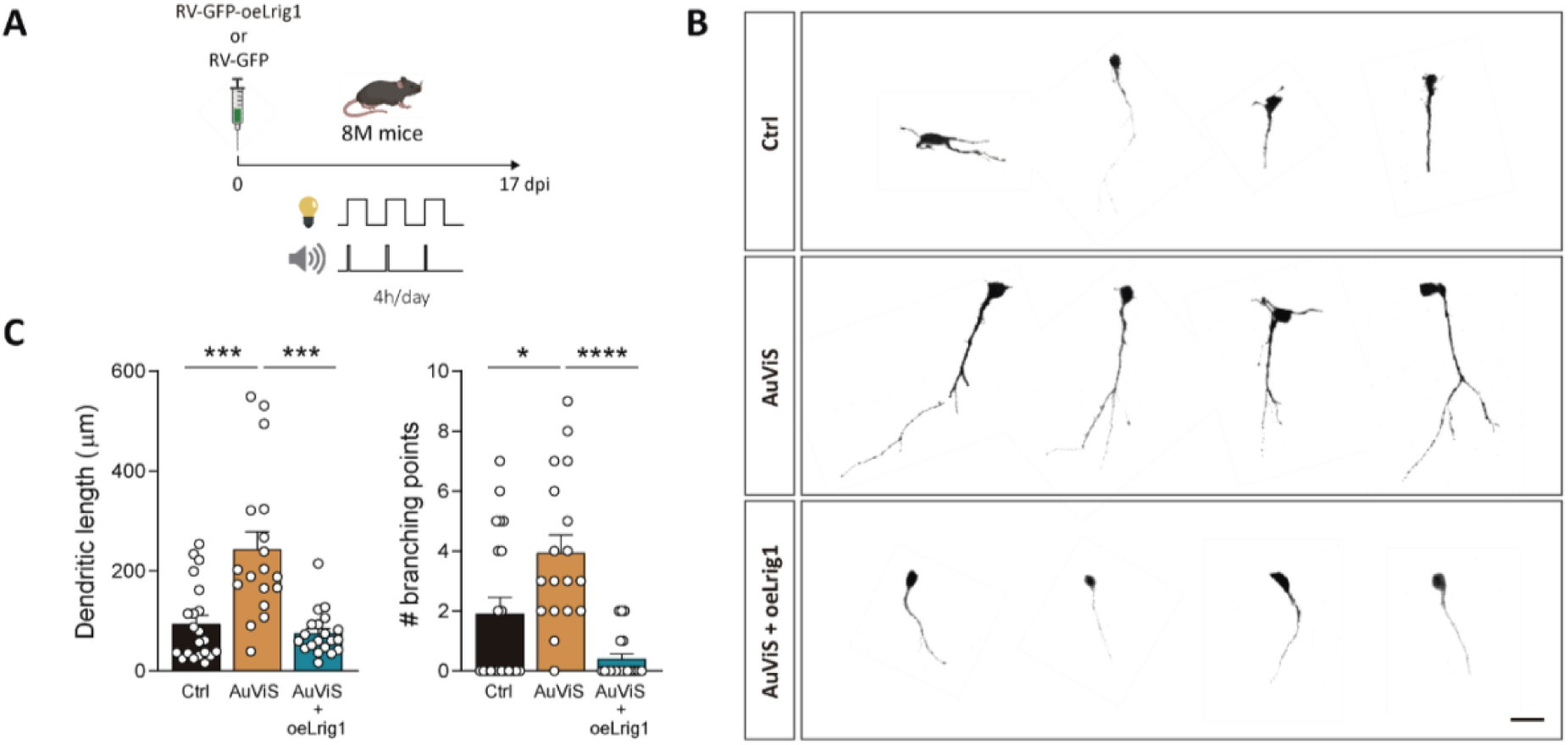
Attenuation of neurotrophin signaling by Lrig1 overexpression precludes the effects of AuViS. (**A**) Experimental design. 8M mice were injected with RV-GFP or RV-GFP-oeLrig1 in the dentate gyrus and exposed to 40 Hz AuViS for 17 days. (**B**) Representative confocal images of 17-dpi GFP-aGCs. Individual neurons have been cropped from the original images. Scale bar, 20 μm. (**C**) Dendritic measurements. (*), and (***) and (****) denote p<0.05, p<0.001 and p<0.0001 after Kruskal-Wallis test followed by Dunn’s post hoc test. Sample sizes (neurons/mice): 21/3 (Ctrl), 18/3 (AuViS) and 20/3 (AuViS + oeLrig1). Bars denote mean ± SEM.

### AuViS enhances proliferation and neuronal differentiation

The experiments described above revealed significant trophic effects of AuViS on aGC development and local circuit remodeling, prompting us to further explore whether AuViS also impacts the quiescence of neural stem cells and the phenotype of their progeny. We assessed the proliferation and differentiation of neural progenitor cells in 11-month-old (11M) mice, a condition whereby neurogenesis displays a minimal rate (*9, 23*). Animals were exposed to AuViS 1 month before and 3 weeks after labeling dividing progenitors using bromodeoxyuridine (BrdU) (**Fig. 5A**). Cell fate was assessed by triple labeling immunofluorescence for BrdU, the neuronal marker NeuN, and the astrocytic marker s100β. AuViS treatment before cell labeling increased the number of BrdU^+^ cells without altering the proportion of neurons and astrocytes, indicating enhanced proliferation of neural progenitors (**Fig. 5B-D**). When maintained after BrdU labeling, AuViS increased neuronal differentiation at the expense of astrocyte production. Together, our results reveal a remarkable potential for 40 Hz multisensory stimulation to enhance neurogenesis in the aging brain at several levels, from increasing the pool of proliferating progenitors and their neuronal differentiation to promoting their growth and integration into the local network.

**Fig. 5.**
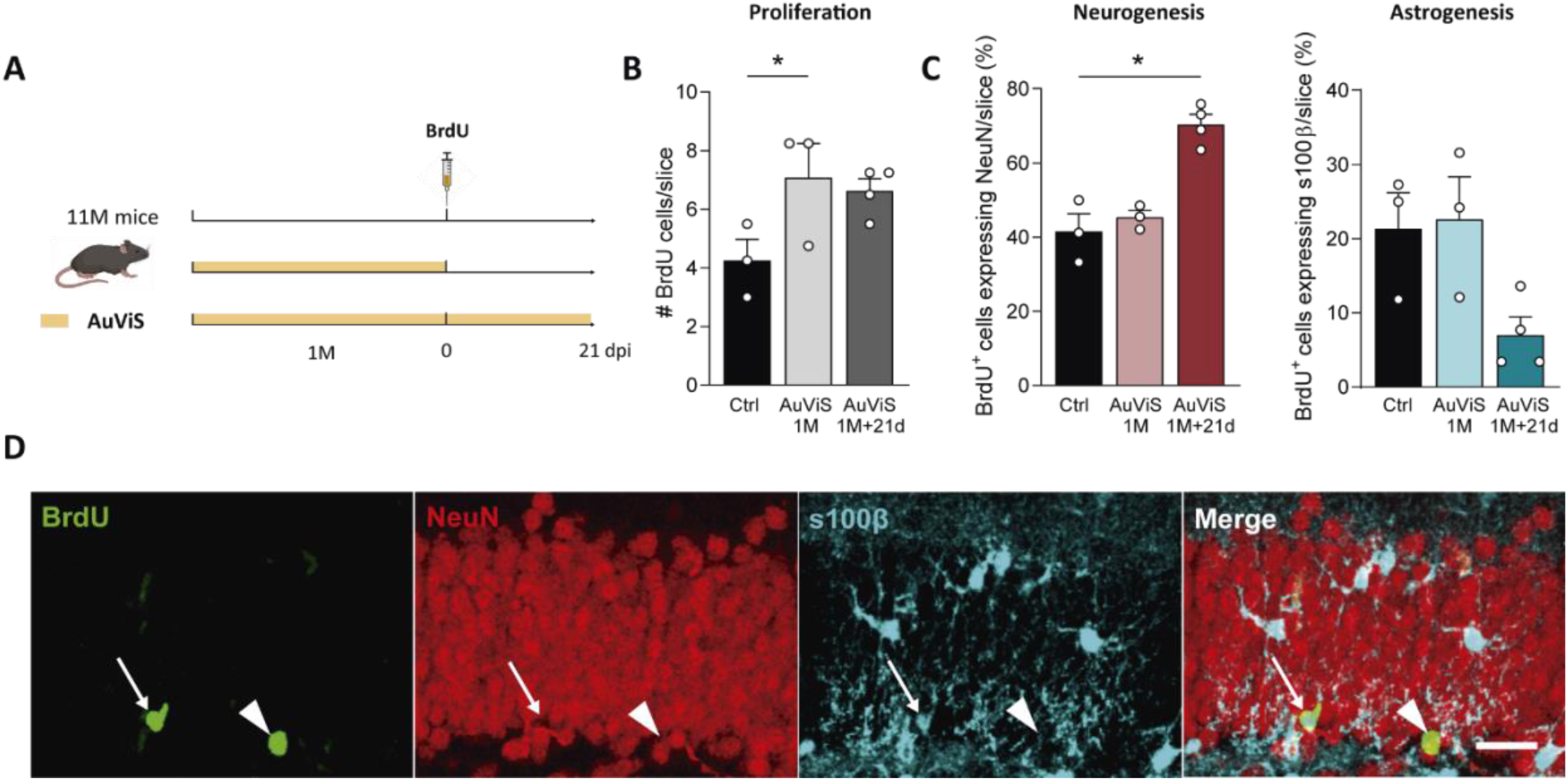
AuViS increases cell proliferation and neurogenesis in the aging brain. (**A**) Eleven-month-old mice were exposed to 40 Hz AuViS or no stimulus and received BrdU as shown in the scheme. (**B**) Total number of labeled cells (BrdU^+^) per slice for all three conditions. (**C**) Fraction of BrdU^+^ cells expressing NeuN or s100β, indicating neurogenesis or astrogenesis. (*) denote *p*<0.05 after Kruskal-Wallis followed by Dunn’s uncorrected post hoc test for multiple comparisons. Sample sizes (mice): 3 (Ctrl), 3 (AuViS 1M) and 4 (AuViS 1M+21d). Bars denote mean ± SEM. (**D**) Representative image showing 21-day-old BrdU^+^ cells (green) co-expressing either NeuN (red, arrowhead) or s100β (blue, arrow). Scale bar, 30 μm.

## Discussion

Gamma-frequency brain oscillations play a critical role in maintaining cognitive function (*24, 25*). Defects in these oscillations have been linked to synaptic dysfunction and cognitive decline (*1, 2*). Specifically, the gamma rhythm abnormalities reported in AD interfere with activity-dependent synaptic potentiation, the cellular substrate for learning and memory (*26*). Even though native gamma oscillations and the state evoked by 40 Hz sensory stimulation are likely distinct neurophysiological phenomena (*27*), non-invasive 40 Hz AuViS promotes Aβ clearance and improve memory in mouse models of AD (*2, 28–31*). These therapeutic effects have been validated across multiple neurological disease models, with one exception where variations in the methodology and experimental design may account for the reported discrepancy (*32*).

One aspect of the overall effects of AuViS that remains unexplored is the activity-dependent plasticity needed to revert the impact of aging and AD on cognitive performance. To address this issue, we sought to leverage the high responsiveness of adult neurogenesis to stimuli that enhance local network activity in the hippocampus of middle-aged animals (*11, 12, 14*). First, we found that AuViS preferentially activated the SGZ of the dentate gyrus (Fig.1A-C), where progenitor cells are located. Given that this region is highly vascularized (*33*), the activation of the glymphatic system exerted by 40 Hz flicker (*34*) could contribute not only to waste elimination but also to a better distribution of glucose, lipids, amino acids and neurotransmitters leading to a positive impact on adult neurogenesis (*35, 36*). AuViS promoted dendritic growth and axonal development, enhancing their morphological features to the point where MFBs resembled those of mature aGCs (Fig. 1D-I) (*12*). Since the gamma component of brain oscillations is still intact in young animals, it was predictable that AuViS would not induce a positive modulation on the dendrites of aGCs in 2M mice (Fig. S6). The decrease in the dendritic tree length observed in 14 (but not 7 or 21) dpi aGCs may be attributable to homeostatic branch pruning in young adult mice (*37*).

Consistent with previous reports, visual stimulation enhances the 40 Hz component of oscillations in the dentate gyrus (Fig. 2) (*2*). Light flickering at 40 Hz but not 8 Hz promoted dendritic growth of aGCs. Different studies in AD models have shown a reduction in Aβ production and neuronal loss at 40 Hz but not at 20 or 80 Hz, highlighting the frequency dependency for the restorative effects of sensory stimulation (*2, 29*). This modulation was also dependent on stimulus duration, as 1 hour of visual stimulation led to a more limited growth of aGC dendrites compared to 4 hours. Auditory stimulation at 40 Hz produced similar changes to those induced by visual flicker. However, when presented individually, neither stimulus was sufficient to elicit presynaptic modifications such as those induced by AuViS (Fig. 1D-I). Accordingly, combined auditory and visual stimulation in the 5XFAD mouse model of AD led to a more extensive reduction of Aβ peptide load across multiple cortical regions, compared to either stimulus presented individually (*28*). The synergy observed in multisensory stimulation likely arises from the simultaneous engagement of multiple neural pathways, enhancing network-wide gamma synchronization, which strengthens inter-regional connectivity (*3*).

AuViS promoted the maturation of neuronal excitability and induced the formation of functional synapses, pointing to its ability to foster the integration of aGCs into the hippocampal circuits (Fig. 3). Effects on synaptic plasticity by 40 Hz stimulation have also been reported in stroke models, where flickering light prevented hippocampal pyramidal cell death with a concomitant increase in LTP induction (*4*), and optogenetic stimulation of interneurons improved cortical connectivity (*38*). Together with the presynaptic effects on the size of MFBs and filopodial growth, our findings demonstrate a remarkable capacity for AuViS to drive mechanisms of synaptic plasticity in the aging brain.

It is likely that decreased neuronal activity, oxidative stress, inflammation, and reduced neurotrophic support all contribute to aging being the primary risk factor for neurodegeneration (*39, 40*). These features could also explain the slow development of aGCs. Activity-dependent neurotrophin release by behavioral stimuli such as running and EE counteract some of the detrimental effects observed both in neurogenesis and neurodegeneration (*11, 12, 41, 42*). We thus tested the hypothesis that gamma stimulation could share some of these mechanisms. Blocking the effects of AuViS by impairing neurotrophin signaling highlighted the important role of trophic factors in neuronal development in the aging brain (Fig. 4). While neurotrophic factors are critical for neurogenesis induced by both exercise and AuViS, multiple mechanisms likely contribute to the complex plasticity observed at the dendritic and axonal compartments (*43*). Indeed, combined treatments of 40 Hz light flickering and running in the 3xTg AD mouse model produced synergistic effects, significantly reducing phosphorylated tau and Aβ peptide load in the hippocampus, suggesting the involvement of distinct, complementary mechanisms (*44*).

AuViS applied before labeling new aGCs increased the rate of cell division, while, when delivered shortly after labeling, it promoted early neuronal differentiation, suggesting that these interventions engage distinct mechanisms at different stages of neurogenesis (Fig. 5A-C). GABAergic interneurons expressing parvalbumin (PV) and vasoactive intestinal peptide (VIP) have been shown to mediate the effects of AuViS (*1, 2, 34*). Specifically, PV interneuron activity can maintain neural stem cell quiescence, but their activation can also promote the development, survival, and integration of aGCs during later stages (*45, 46*). In contrast, the firing of mossy cells is known to activate quiescent stem cells and could contribute to the neurogenic effects observed here (*47*).

Due to its broad accessibility, gamma multisensory stimulation holds significant potential for providing therapeutic benefits to patients suffering from neurological disorders, including non-degenerative diseases such as epilepsy (*48*). Several studies have shown that this treatment is safe and feasible in humans (*49–51*). A deeper understanding of the mechanisms underlying overall cognitive improvement could substantiate AuViS as a therapeutic approach for neurological diseases that are increasingly prevalent and currently lack effective treatments (*52, 53*).

## Materials and Methods

### Mice and surgery

C57BL/6J (2, 8 and 12 months of age) and genetically modified (8 months of age) male and female mice were housed at two to four animals per cage in standard conditions. Middle-aged mice were selected because, beyond this age, there is a strong decline in hippocampal neurogenesis that precludes the study of labeled neurons (*9, 11*). *Ascl1^CreERT2^ (Ascl1^tm1(Cre/ERT2)Jejo^/J)* mice (*54*) were crossed to a *CAG^floxStoptdTomato^* (Ai14) *[B6;129S6-Gt(ROSA)26Sor^tm14(CAG-tdTomato)Hze^/J]* conditional reporter line (*55*) to generate *Ascl1^CreERT2^;CAG^floxStopTomato^*mice, which can be used to reliably target adult-born aGCs (*56*). Tamoxifen (TAM) administration (120 mg/kg, once a day for 3 days) was carried out in 8M mice to achieve indelible expression of tdTomato (Tom) in the progeny of Ascl1^+^ progenitor cells. Two to three days before TAM induction or retroviral injection, mice were exposed to running wheels (one for every two mice) to maximize the number of labeled adult-born aGCs. Mice were anesthetized (150 µg ketamine/15 µg xylazine in 10 µl saline per gram), and retrovirus was infused into the septal region of the right dentate gyrus (1.5 µl at 0.15 µl/min) using sterile calibrated microcapillary pipettes through stereotaxic surgery; coordinates from bregma (in mm): −2 anteroposterior, −1.5 lateral, and −1.9 ventral. In order to perform morphological analysis, brains were fixed after perfusion and sections were prepared at the indicated times for confocal imaging (*11*). Only aGCs from the septal dentate gyrus were included in the analysis, corresponding to sections localized from −0.96 to −2.30 mm from the bregma, according to the mouse brain atlas (*57*). Experimental protocols were approved by the Institutional Animal Care and Use Committee of Fundacion Instituto Leloir, according to the Principles for Biomedical Research involving animals of the Council for International Organizations for Medical Sciences and provisions stated in the Guide for the Care and Use of Laboratory Animals.

### Bromodeoxyuridine labeling

To assess the effects of AuViS on proliferation, bromodeoxyuridine was dissolved in 0.9% NaCl and was delivered to 12M mice intraperitoneally at a dose of 50µg/g to label dividing cells. Mice received two injections per day over 3 days and were sacrificed 21 days later.

### Auditory and visual stimulation protocols

#### Visual stimulation protocol

To perform visual stimulation, we used white LEDs, which include a combination of visible wavelengths (390–700 nm). Specifically, we used LEDs that have a correlated color temperature (CCT) of 4,000 K at intensities between 100 and 480 lux. After stereotaxic injection, mice were placed in a standard cage with three sides covered with black plastic and one transparent side facing a LED strip. This configuration was used to ensure that the LED strip is the main light source and to visually separate the animals (**Fig. S1**). Mice were exposed to one of two stimulation conditions: 8 Hz or 40 Hz stimulation (50 % working cycle) for 4h/day for 17 days (**Fig. S1**). In **Figure S3**, animals were exposed to 40 Hz flicker for 1h/day for 21 days. Intensity was set at ∼480 lux at the side closest to the lights as previously reported (*58*). Control mice were left in standard cages.

#### Auditory stimulation protocol

Animals were placed in the same cages used for visual stimulation. Speakers (Genius, SP-HF500A) were placed between cages (**Figure S1**), and mice were exposed to tones (10 kHz, 1 ms, 60 dB) at 40 Hz for 4h/day for 17 days (*58*).

#### Audiovisual stimulation protocol

Animals were simultaneously exposed to synchronous visual flicker and auditory tone trains. Mice were exposed to one of two stimulations: 40 Hz (AuViS) or random (50% working cycle, random intervals) stimulation for 4h/day for 17 days. To assess the expression of the immediate early gene (IEG) ARC, mice were exposed to AuViS for 2 h and brains were collected 1 hour later (**Fig. 2**). For slice physiology recordings, mice were exposed to AuViS for 17 or 24 days (**Fig. S5 and Fig. 3**). Young animals were exposed to AuViS for 7, 14 and 21 days (**Fig. S6**). To assess proliferation, 11-month-old mice were exposed to AuViS for 1M or 1M plus 21 days (**Fig. 5**).

### In vivo electrophysiology

#### Drives

Tetrodes (∼25-um in diameter) were constructed from four strands of tungsten wire (99.95% tungsten, 12.5 um in diameter, California Fine Wire Company, USA), bound together by twisting and melting their insulation with hot air (approximately 110° C). Five tetrodes were loaded and glued into a nested assembly of stainless-steel tubes and mounted into the microdrive (Axona Limited, UK). The tetrodes exited the microdrive through a guide cannula in an approximately rectangular arrangement (approximate spacing 400-µm) and every tetrode could be moved independently via a drive screw (160 µm per turn). Each tetrode was cut flat and its tip was gold-plated (gold plating solution, Neuralynx, MT, USA) to reduce the 1 KHz impedance of individual wires to 0.2–0.4 MOhm (Nano Z, White Matter, WA, USA) (*59*).

#### Surgery for in vivo experiments

The surgeries to chronically implant the drives were conducted under ketamine-xylazine anesthesia (150 μg ketamine/15 μg xylazine in 10 μl saline/g). A circular opening (approximately 1 mm in diameter) was made in the skull above the right dorsal hippocampus. The center of the craniotomy was 1.5 mm lateral to the midline and 2 mm posterior to bregma. Next, 7 jeweler screws were inserted in holes drilled in the skull around the craniotomy. After the removal of the dura, the tetrode tips were lowered 1 mm below the brain surface. The craniotomy was filled with a biocompatible transparent gel Neuroseal obtained by combining equal parts of 0.5 % sodium alginate and 10 % calcium chloride, both previously dissolved in distilled water and stored separately. The skull, the screws and the base of the opto-microdrives were then covered with autocrystal acrylic, dental cement. After surgery, animals were left to recover for 3-5 days.

#### Lowering of tetrodes

During a period of 2 – 3 weeks after recovery from surgery, tetrodes were progressively lowered toward their target subfields in the hippocampus. Dentate spikes and granule cell activity, visualized with Cheetah (Neuralynx, Inc., MT, USA), were the main guides to position the electrodes in the dentate gyrus.

#### Recordings

Mice were connected to the recording cable and put in a cage with a wire mesh around it, which ensured that light would pass while serving as a Faraday cage. LEDs were distributed around the cage outside the mesh and activated according to the desired condition through an Arduino board controlling the power input to the LEDS through a MOSFET transistor.

#### Local field potential analysis

All analyses were performed using MATLAB scripts. Power spectral densities were obtained using the pwelch() function. To compare the power spectral density at 40 Hz between two 5-minute sessions, sessions were divided into non-overlapping 10 s windows and the power spectral density at 40 Hz for each window was obtained. The distribution of power spectral densities for both groups of windows was compared using a Mann-Whitney test.

#### Brain processing for histology

Electrodes were not moved after the final recording session. Subjects underwent an intraperitoneal injection of ketamine-xylazine anesthesia, were perfused intracardially with 100 ml of heparinized saline solution, and then with 100 ml of 4 % paraformaldehyde. Three hours after perfusion, tetrodes were raised, the brain was stored in 4% paraformaldehyde overnight, and then put in sucrose solution (30 %) until it precipitated (around 48 h). Frozen coronal sections (40 μm) were cut with a microtome (Leica, Germany) and dorsal hippocampal sections were mounted on glass slides. After around 20-minute drying, they were covered with PVA-DABCO mounting media and coverslips. The fluorescence microscope Zeiss Axio Examiner.D1 were used to obtain digitalized images.

### Immunofluorescence

Immunostaining was done on 60-µm free-floating coronal sections. Antibodies were applied in tris-buffered saline (TBS) with 3% donkey serum and 0.25% Triton X-100. Single or double labeling immunofluorescence was performed using the following primary antibodies: GFP (rabbit polyclonal; 1:500; Invitrogen), GFP (chicken polyclonal; 1:500; Aves), Arc (rabbit polyclonal; 1:500; Synaptic Systems), NeuN (mouse monoclonal; 1:50; a gift from F.H. Gage, Salk Institute for Biological Studies, La Jolla, CA), S100b (rabbit monoclonal; 1:500; abcam), BrdU (rat monoclonal; 1: 250; Abcam), RFP (rabbit polyclonal; 1:500; Rockland Immunochemicals) and RFP (goat polyclonal; 1:500; Rockland Immunochemicals). The following corresponding secondary antibodies were used: donkey anti-rabbit Cy3, donkey anti-rabbit Cy5, donkey anti-mouse Cy5, donkey anti-goat Cy3 and donkey anti-chicken Cy2 (1:250; Jackson ImmunoResearch Laboratories). For BrdU detection, DNA was denatured with 50% formamide in 2X SSC buffer at 65°C for 1 h, washed twice in 2XSSC for 10 min, incubated in 2N HCl at 37°C for 30 min, and washed in 0.1 M boric acid, pH 8.5, for 10 min. The steps of a standard protocol were then followed.

### Confocal microscopy

Images were acquired using confocal microscopes. For analysis of Arc expression, images were acquired using a Zeiss LSM 880 confocal microscope (40x objective; NA, 1.3; oil immersion), and colocalization was assessed in Z stacks using multiple planes for each cell. For dendritic length measurements, images were acquired using a Zeiss LSM 510-META confocal microscope (40x objective; NA 1.3; oil-immersion) from 60-µm thick sections taking Z stacks including 35–50 optical slices, airy unit = 1 at 0.8-µm intervals (*12*). Dendritic length was then measured using the LSM Image Browser software from projections of three-dimensional reconstructions onto a single plane in aGCs expressing GFP. Images of GFP-labeled MFBs in the CA3 region were acquired at 0.4-μm intervals using a Zeiss LSM 880 confocal microscope (63x objective; NA 1.4; oil-immersion) and a digital zoom of 6. Area and number of extensions of mossy fiber boutons were analyzed from projections of three-dimensional reconstructions onto a single plane. Mossy fiber boutons that fit the following criteria were selected for quantification: (i) the diameter of the bouton was >3-fold larger than the diameter of the fiber, (ii) the bouton was connected to the mossy fiber on at least one end (*16*). Extensions were identified as protrusions arising from large mossy terminals (1 μm < length < 20 μm) (*15*). Filopodial extensions were measured by counting the number of protrusions per terminal. For image capture and analysis of morphological properties, all experimental groups under study were blind to the operator.

### Hippocampal slice electrophysiology

#### Slice Preparation

Ascl1^CreERT2^;CAG^floxStop-tdTomato^ 8M mice were anesthetized, and transverse slices were prepared at 17 or 24 days after TAM induction as indicated (**Fig. 3 and Fig. S5**) (*11, 46*). Briefly, brains were removed into a chilled solution containing (in mM): 110 choline-Cl, 2.5 KCl, 2.0 NaH_2_PO_4_, 25 NaHCO_3_, 0.5 CaCl_2_, 7 MgCl_2_, 20 dextrose, 1.3 Na^+^-ascorbate, 3.1 Na^+^-pyruvate, and 4 kynurenic acid. Coronal slices (400-µm thick) from the septal pole containing both hippocampi were cut with a vibratome and transferred to a chamber containing (in mM): 125 NaCl, 2.5 KCl, 2 NaH_2_PO_4_, 25 NaHCO_3_, 2 CaCl_2_, 1.3 MgCl_2_, 1.3 Na^+^-ascorbate, 3.1 Na^+^-pyruvate, and 10 dextrose (315 mOsm). Slices were bubbled with 95% O_2_/5% CO_2_ and maintained at 30°C for >45 min before experiments started.

#### Recordings

Whole-cell recordings were performed using microelectrodes (4–6 MΩ) filled with (in mM): 150 K-gluconate, 1 NaCl, 4 MgCl_2_, 0.1 EGTA, 10 HEPES, 4 ATP-Tris, 0.3 GTP-Tris, and 10 phosphocreatine. Criteria to include cells in the analysis were visual confirmation of tdTomato in the pipette tip, attachment of the labeled soma to the pipette when suction was performed, and absolute leak current <100 pA at holding potential (V_h_). Spontaneous EPSCs were recorded in voltage clamp at −70 mV. Input resistance was assessed by the application of voltage steps of −10 mV in voltage-clamp mode, and spiking was evoked by the injection of current steps (10 pA) in current-clamp configuration after taking the membrane potential to −70 mV. All recordings were performed at room temperature (23 ± 2 °C), digitized, and acquired at 10 KHz on a personal computer. Detection and analysis of spontaneous EPSCs were done using a dedicated software package.

### Statistical analysis

Statistics used throughout the paper are described in the figure legends and in the text. Unless otherwise specified, data are presented as mean ± SEM. Normality was assessed using the Shapiro-Wilks test, D’Agostino-Pearson omnibus test, and Kolmogorov-Smirnov test, with a *p*<0.05. When data met normality tests (Gaussian distribution and equal variance), unpaired *t*-test with Welch’s correction or ANOVA with Bonferroni’s post-hoc test were used as indicated. In cases that did not meet normality, nonparametric tests were used as follows: Mann-Whitney test for independent comparisons, and Kruskal-Wallis test for multiple comparisons.

## Acknowledgments

We thank members of the A.F.S. and E.K. labs for insightful discussions, Agostina Miranda and Juan Simón Serrangeli for data collection (Figs. 1C, 2H and S6). M.F.T., E.K. and A.F.S. are investigators at the Consejo Nacional de Investigaciones Cientificas y Tecnicas (CONICET). M.H., M.M. and S.B. were supported by CONICET fellowships.

## Author contributions

Conceptualization: MFT, EK, AFS

Methodology: SB, IGS

Investigation: MFT, MH, AAA, MM

Visualization: MFT, MH, EK, AFS

Funding acquisition: MFT, EK, AFS

Project administration: MFT, AFS

Supervision: MFT, EK, AFS

Writing – original draft: MFT, MH, EK, AFS

## Data and materials availability

All data are available in the main text or the supplementary materials.

