## Supplemental Figures for "Audiovisual gamma stimulation enhances hippocampal neurogenesis and neural circuit plasticity in aging mice"

**This file includes:**

Figs. S1 to S6

**Fig. S1**

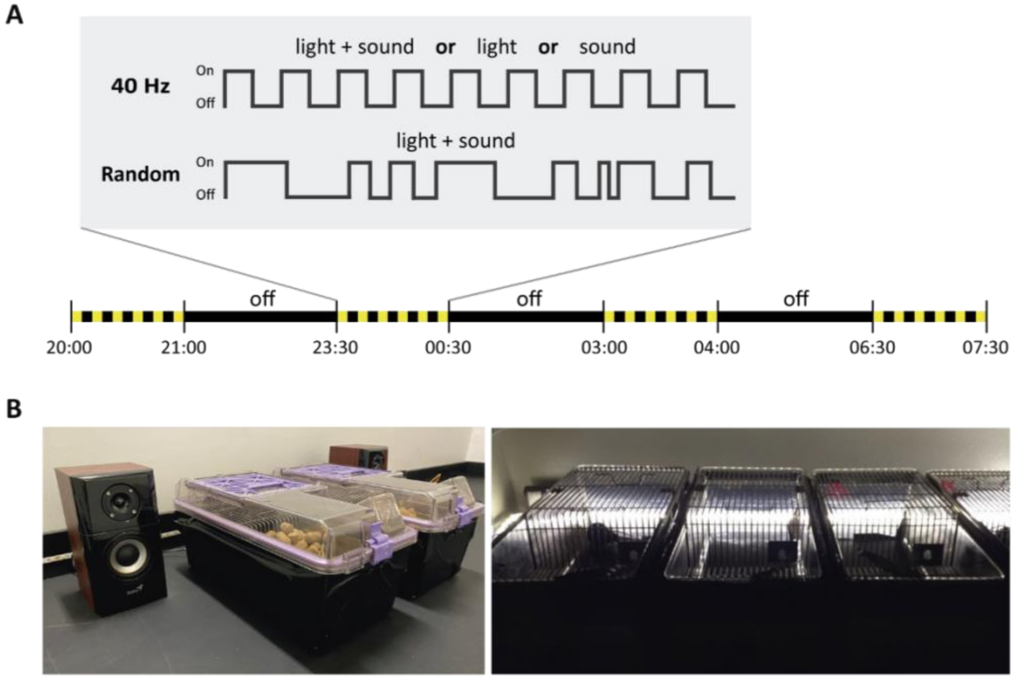


**AuViS stimulation setup.** (**A**) Schematic representation of 40 Hz and random stimuli patterns. Stimuli consist of synchronous flashing lights (4,000 K, 50 % working cycle, 480 lux) and tones (10 kHz, 1 ms, 60 dB), delivered in four 1 h-intervals separated by 2.5 hours of darkness and repeated daily for the indicated durations. (**B**) The flicker setup consists of LED strips mounted along a wall and two speakers distributed to the left and right sides of the cages. Cages are covered on three sides, with the transparent side oriented towards the LEDs. Further details in the **Methods** section.

**Fig. S2**

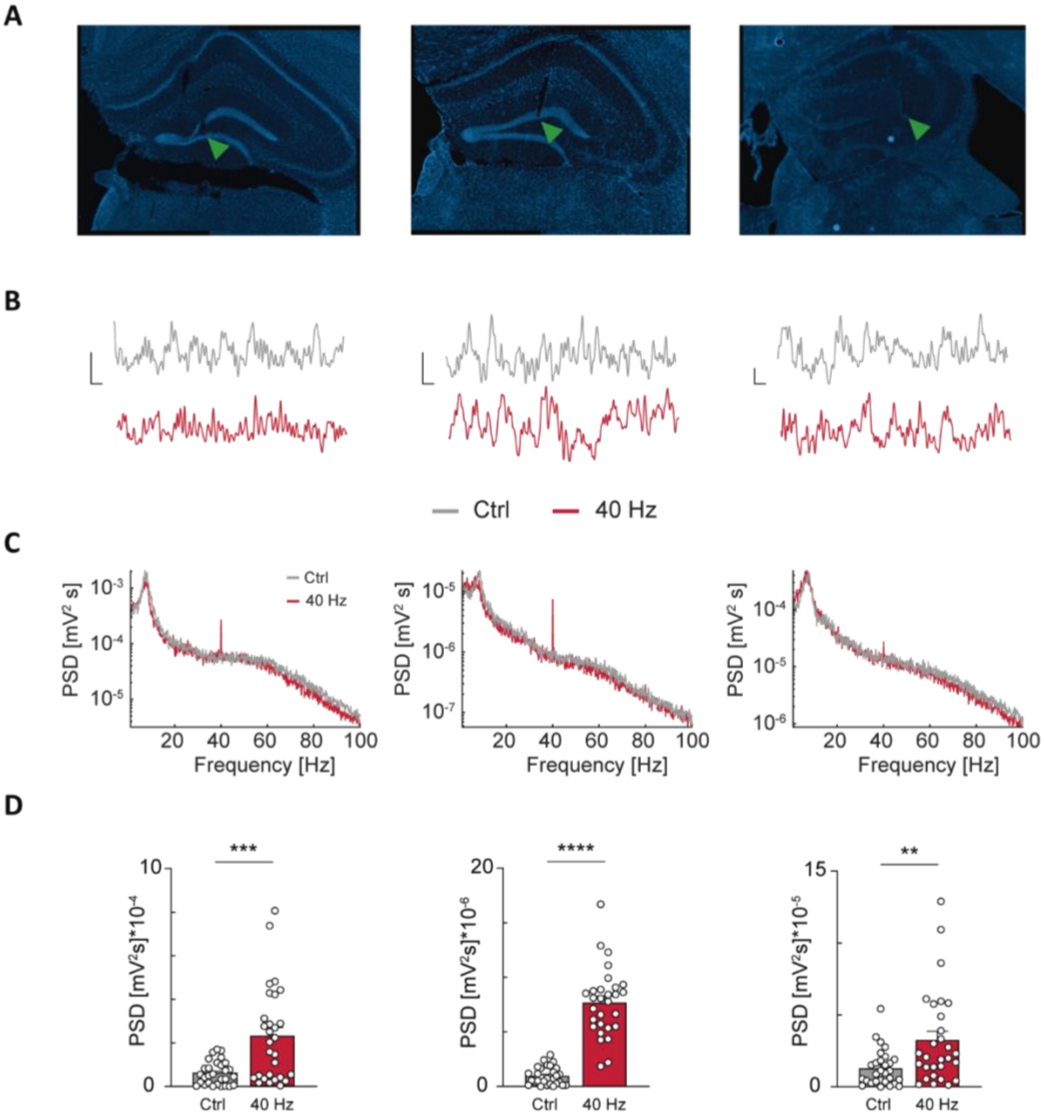


**Visual stimulation at 40 Hz modulates the local field potential in the dentate gyrus.** (**A**) Fluorescent images (DAPI) displaying the traces of tetrodes (one per mouse) targeted to the granule cell layer (green arrowhead: tetrode tip). (**B**) Representative example trace of 1 s raw local field potential during flickering (40 Hz, red) or lights-off (Ctrl, grey) control conditions. Scale bars (left to right): 500 μV, 0.1 s; 50 μV, 0.1 s; 100 μV, 0.1 s. (**C**) Corresponding power spectral densities for 5 min of flickering (40 Hz) or lights-off (Ctrl). (**D**) Power spectral density at 40 Hz for consecutive non-overlapping 10 s windows of data. (**), (***) and (****) denote $p<0.01$, $p<0.001$ and $p<0.0001$ after Mann–Whitney test. Bars denote mean ± SEM.

**Fig. S3**

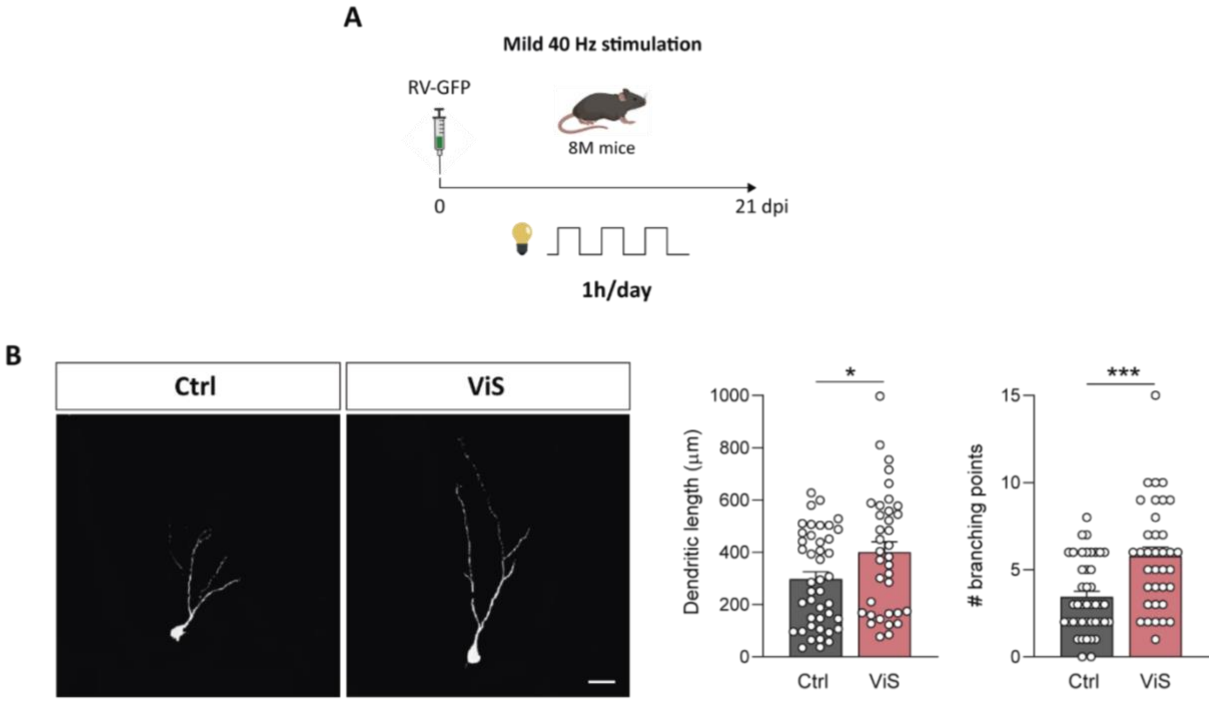


**Effects of mild visual stimulation on aGCs in 8M mice.** (**A**) RV-GFP injection was followed by 21 days to 40 Hz AuViS (1 h/day). (**B**) Left: Representative confocal images of 21-dpi GFP-aGCs. Scale bar: 20 µm. Right: Analysis of dendritic morphology. (*) and (***) denote $p$<0.05 and $p$<0.001 after Mann-Whitney test. Sample sizes (neurons/mice): 43/4 (Ctrl) and 36/4 (ViS). Bars denote mean ± SEM.

**Fig. S4**

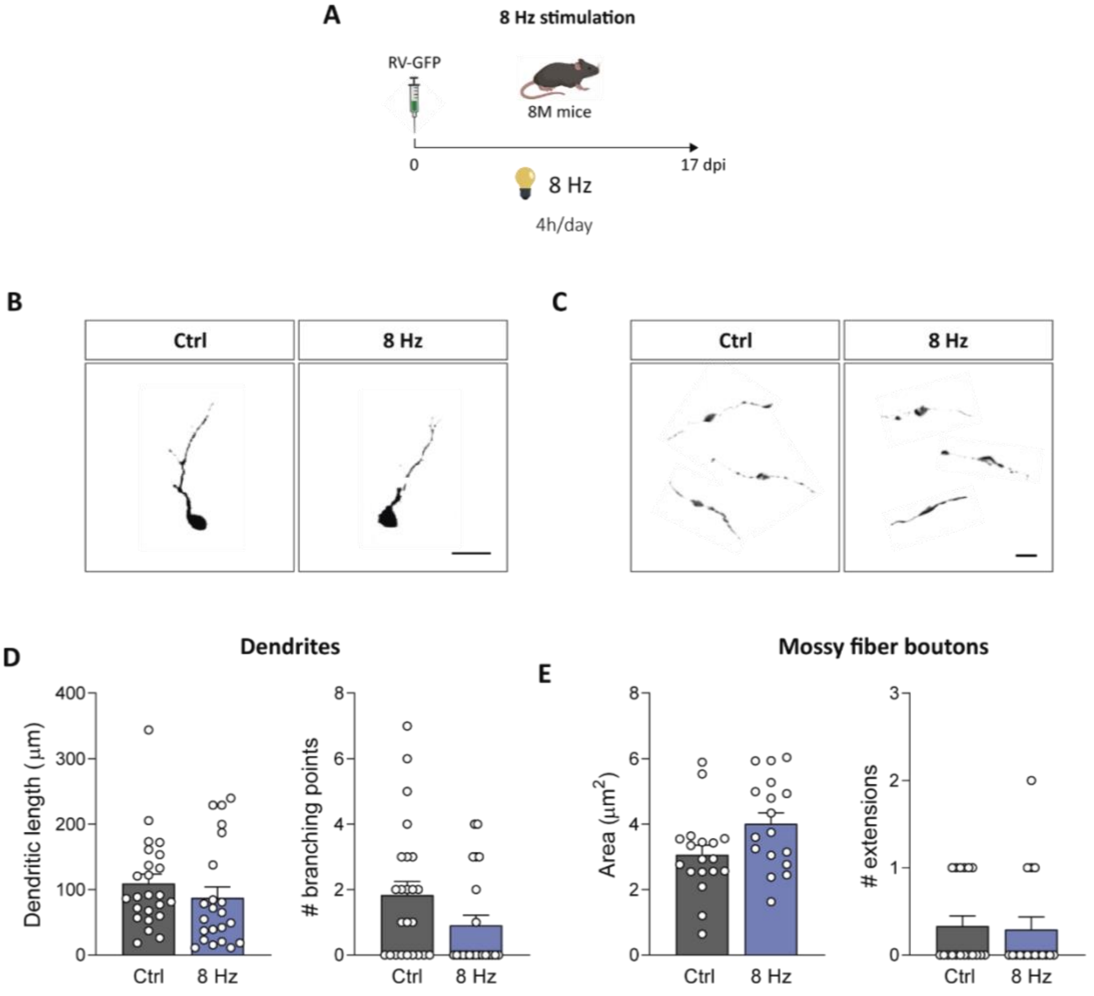


**Visual stimulation at 8 Hz does not influence aGC development.** (**A**) RV-GFP injection was followed by 17 days of 8 Hz AuViS. (**B**) Representative confocal images of 17-dpi GFP-aGCs. Individual neurons have been cropped from original images. Scale bar, 20 μm. (**C**) Representative images of 17-dpi MFBs in CA3 for the two groups. Scale bar, 5 μm. (**D**) Dendritic measurements. Sample sizes (neurons/mice): 24/3 (Ctrl) and 22/3 (8 Hz). (**E**) MFB architecture. Sample sizes (neurons/mice): 18/3 (Ctrl) and 17/3 (8 Hz). Bars denote mean ± SEM.

**Fig. S5**

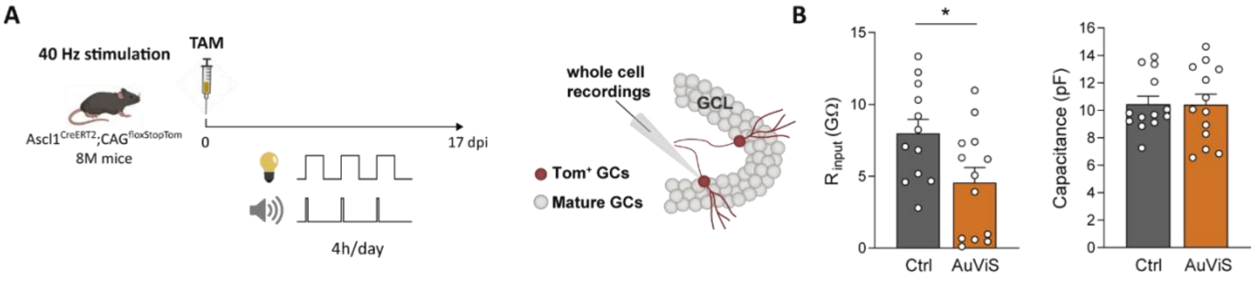


**AuViS modifies the intrinsic properties of 17 dpi aGCs.** (**A**) *Ascl1*$^{CreERT2}$*;CAG*$^{floxStoptdTomato}$ 8M mice received TAM to label new aGCs. Acute slices were prepared 17 days after 40 Hz AuViS or control treatments to assess intrinsic properties and connectivity using whole-cell recordings in tdTomato$^{+}$-aGCs. (**B**) Input resistance and membrane capacitance. Sample sizes (neurons/mice): 12/3 (Ctrl) and 13/2 (AuViS). (*) denotes $p$<0.05 after Mann–Whitney test. Bars denote mean ± SEM.

**Fig. S6**

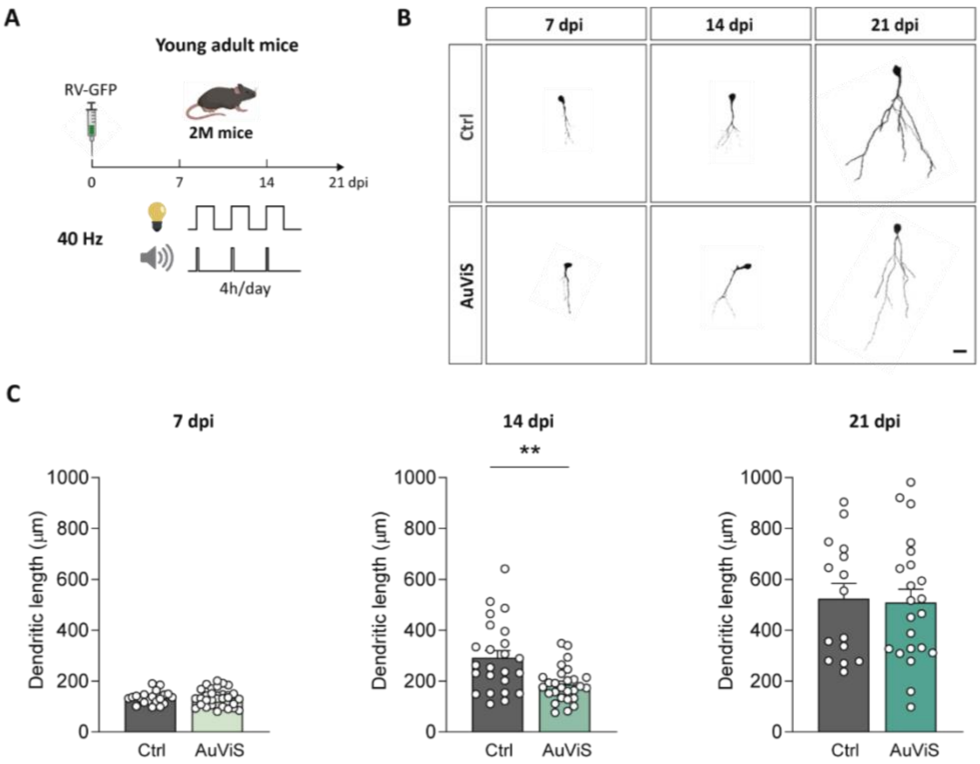


**AuViS does not accelerate GCs development in young mice.** (**A**) Two-month-old mice were injected with a RV-GFP and exposed to 40 Hz AuViS for 7, 14 or 21 days. (**B**) Representative confocal images of GFP-aGCs at different developmental ages. Scale bar, 20 µm. (**C**) Dendritic measurements. (**) denote $p<0.01$ after t test. Sample sizes (neurons/mice) for 7 dpi: 20/3 (Ctrl) and 31/4 (AuViS); 14 dpi: 24/4 (Ctrl) and 27/4 (AuViS); 21 dpi: 15/3 (Ctrl) and 22/4 (AuViS). Bars denote mean ± SEM.
